# Beta power modulation supports micro-consolidation of implicit motor sequences

**DOI:** 10.64898/2026.07.26.740849

**Authors:** Joshua Hendrikse, Nermin Aljehany, Juliet Hosler, Ella Saffery, Emily Brooks, James P. Coxon

## Abstract

The consolidation of novel motor skills has traditionally been investigated over the timescale of hours-days following practice. However, a growing body of evidence has demonstrated that consolidation of skills occurs more rapidly across a scale of seconds – a process termed ‘micro-consolidation’. Micro-consolidation of explicitly cued motor sequences is supported by frontoparietal beta oscillations. Whether beta power modulation is a common mechanism supporting rapid consolidation of other forms of skill learning, such as implicit sequence learning, is yet to be elucidated. 72 healthy adults aged 18-35 (55% female) completed a serial reaction time task with concurrent electroencephalography recording. In line with our previous work, we show that the early ‘fast’ learning on an implicit sequence task is primarily expressed as micro-offline gains during brief rest periods between periods of practice. Beta power was modulated as a function of active practice vs rest epochs (i.e., event-related synchronisation/desynchronisation), and here we demonstrate that micro-offline gains are associated with this modulation of beta power during the rest epochs. This relationship was specific to the beta frequency, and was not observed across either mu or gamma bands. Overall, in line with seminal work implicating beta in early explicit motor learning, our results indicate that beta is a shared neurophysiological signature of micro-consolidation of implicit sequences.

**Key points summary:**

- Recent evidence demonstrates rapid consolidation of motor skills over seconds, termed micro-consolidation.
- Seminal work has implicated beta oscillations in early explicit motor learning, though whether beta supports rapid consolidation of other forms of skill learning is unclear.
- Using electroencephalography, we demonstrate that beta modulation is a neurophysiological signature of micro-consolidation of implicitly learnt motor sequences.
- Lower beta at rest is associated with a higher degree of micro-consolidation.
- Our findings shed critical new insight into the neurophysiological mechanisms mediating rapid consolidation of implicit motor skills during early ‘fast’ learning.

## Introduction

The expression of complex motor skills stems from motor skill learning – a process involving skill acquisition, ‘offline’ consolidation during rest, and retention (Dayan & Cohen, 2011). Motor skill learning can be broadly conceptualised across two key phases – an initial ‘fast’ learning phase associated with rapid skill improvement, followed by a ‘slow’ learning phase associated with comparatively incremental gains in skill and performance (Dayan & Cohen, 2011; Doyon & Benali, 2005; Ungerleider et al., 2002). A growing body of research has demonstrated that during initial fast learning, consolidation commences rapidly across a scale of seconds during brief rest periods between bouts of practice– a process referred to as micro-consolidation (Bönstrup et al., 2019; Brooks et al., 2026). The micro-consolidation phenomenon has primarily been investigated during explicit skill learning requiring execution of a cued motor sequence, and is linked to frontoparietal beta oscillations (Bönstrup et al., 2019; Buch et al., 2021; Jacobacci et al., 2020) and neural replay across cortico-hippocampal networks (Buch et al., 2021; Chen et al., 2024; Griffin et al., 2025; Sjøgård et al., 2025).

Recent work has demonstrated that micro-consolidation generalises beyond explicit sequence learning to *implicit* skill learning (Brooks et al., 2024, 2026; Griffin et al., 2025), which develops without conscious awareness. However, relative to explicit skill learning, the neural mechanisms underlying micro-consolidation of implicit learning in humans are unclear. This is a critical gap in knowledge given the central role of implicit learning processes in the development of autonomous motor abilities (Dayan & Cohen, 2011; Doyon & Benali, 2005), and evidence of disrupted implicit learning processes in movement disorders such as Parkinson’s disease (Siegert et al., 2006). An improved understanding of the neurophysiological processes underlying early stages of skill learning could inform more targeted, effective intervention.

A considerable body of research has implicated beta (13–30 Hz) oscillations in sensorimotor control and skill learning (Engel & Fries, 2010; Tan et al., 2014; Torrecillos et al., 2018), including the acquisition and retention of implicit motor skills (Pollok et al., 2014). During early learning, a reduction in beta power occurs during action preparation and execution – a phenomenon known as event-related desynchronisation (ERD) (Pfurtscheller et al., 1997; Pollok et al., 2014). A transient rebound in beta power is then observed upon action termination (event-related synchronisation, ERS), which has been linked to error monitoring and adaptation (Koelewijn et al., 2008; Tan et al., 2014; Torrecillos et al., 2018; Wu et al., 2025). ERD/S are associated with gamma-aminobutyric acid (GABA)-mediated inhibition (Gaetz et al., 2011; Hall et al., 2011; Muthukumaraswamy et al., 2013), which plays a critical role in ‘gating’ the synaptic plasticity associated with skill learning (Hendrikse et al., 2025; Stagg et al., 2011). Notably, Bonstrup et al. (2019) demonstrated that greater micro-consolidation of an explicitly cued motor sequence was associated with reduced beta power during the brief rest epochs, plausibly reflective of disinhibition processes mediating early learning. However, whether beta power modulation (e.g., beta ERD/S) is also implicated in micro-consolidation of implicit learning is yet to be established.

The present study investigated beta power modulation during micro-consolidation of implicit motor sequences. It was hypothesised that early learning of an implicit motor sequence would be primarily attributable to gains in skill during brief offline periods of rest (i.e. ‘micro-offline’gains). It was also expected that higher micro-offline gains would be associated with reduced beta power measured during brief rest periods between practice blocks. To examine the specificity of these effects, we also explored associations across mu and gamma power bands, given previous work establishing a role of these oscillations in motor performance (Hussain et al., 2021; Nowak et al., 2018).

## Materials and Methods

### Ethical Approval

This study was approved by the Monash University Human Research Ethics Committee (project # 27724), and all participants provided their written informed consent. The study conformed to the standards set by the Declaration of Helsinki, except for registration in a database. Participants were remunerated $30 for their participation.

### Participants

72 right-handed healthy adults (M_age_ = 22.26 ± 3.67 SD, range = 18-35 years, 55% female) without history of neurological or psychiatric illness, use of psychoactive medication, or previous participation in research involving motor sequence learning tasks were recruited through convenience sampling methods. This sample size was supported by an *a priori* power analysis which indicated N = 67 to detect a medium-large effect size, informed by the seminal work on micro-consolidation by Bönstrup et al. (2019) reporting associations between beta power and micro-offline learning of an explicit motor sequence.

### Design

This study employed a cross-sectional, within-subjects design. Participants attended a single session wherein EEG data was obtained before and during practice of a motor sequence learning task.

### Procedure

#### EEG

64 channels of continuous EEG data were recorded at a 1000 Hz sampling rate (low cut off: DC; high cut-off: 280 Hz) using an Ag/AgCl electrode cap (ActiCHamp Plus, Brain Products, Germany), BrainVision Recorder software (Brain Products, Germany), and an ActiCHamp amplifier. Electrodes were arranged in accordance with the international 10-20 montage with channel FCz as the online reference, and impedances were kept below 5 kΩ. Three minutes of resting-state data were first recorded, with participants instructed to focus on a central fixation cross presented in black on a white background and to “let their mind go blank and not think about anything in particular”. EEG data were then acquired as participants completed a serial reaction time task (SRTT).

#### Serial Reaction Time Task (SRTT)

In this task, participants were required to respond to visual cues presented on a computer monitor using a button box and the fingers of their non-dominant left hand (see Figure 1a). Four white circles were displayed horizontally and on each trial participants were instructed to respond as quickly and accurately as possible to the circle that was presented in black by pressing the corresponding button on the button box. Unknown to the participant, the visual cues were presented in a 12-item repeating sequence (2-3-1-4-3-2-4-1-3-4-2-1). The sequence was designed to ensure that each element (1 to 4) occurred 3 times in the sequence, and no element occurred twice in a row. The task included a total of 24 blocks, where each included a 10 second practice period in which participants were instructed to respond to visual cues, and a 10 second rest period (i.e., 480 seconds of total practice). After completing the task, the participant’s level of explicit sequence awareness was assessed. Participants who were able to successfully reproduce at least five consecutive sequence items were deemed to have developed explicit sequence awareness, and were excluded from subsequent analysis (n = 7).

**Figure 1.**
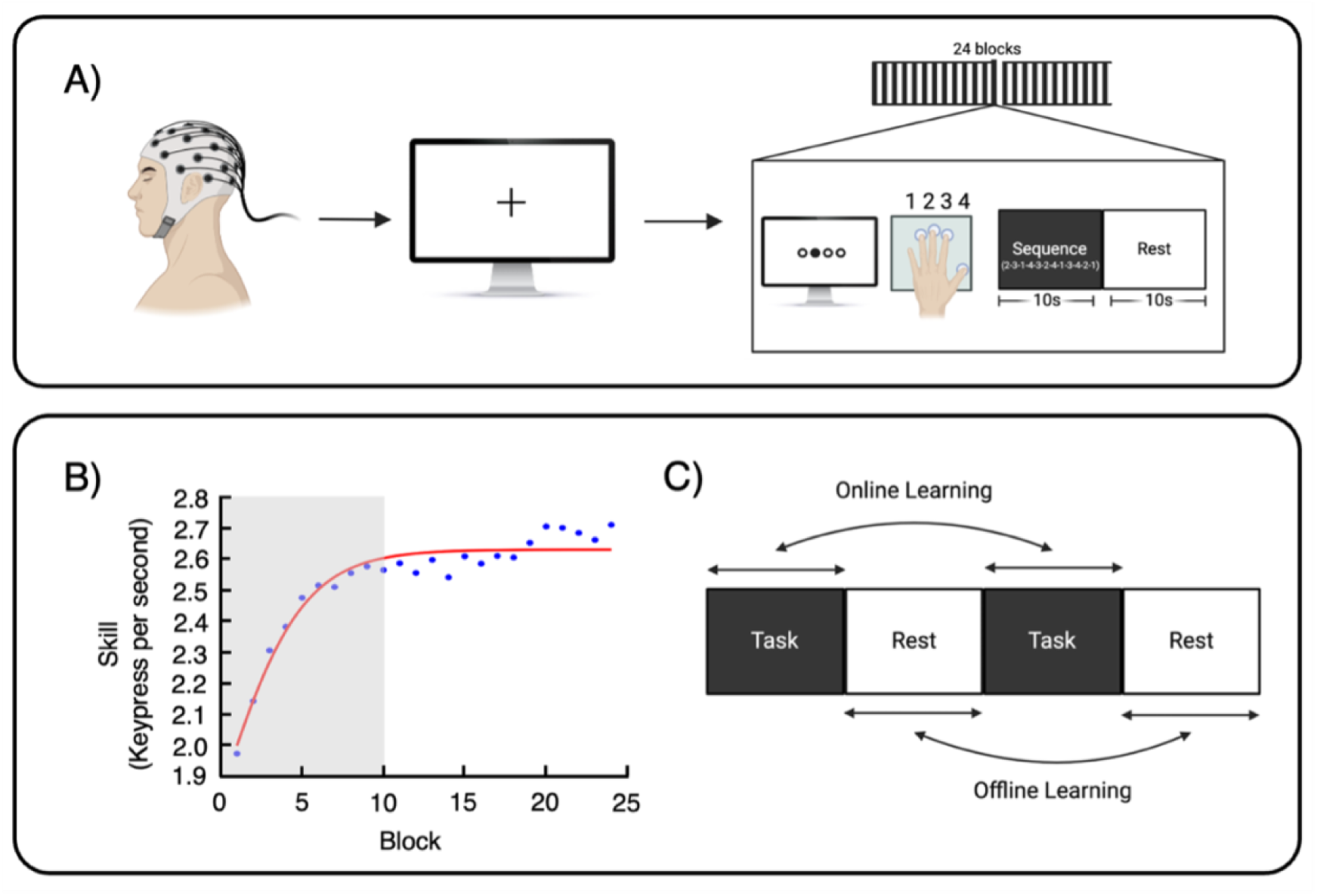
*A,* Overview of the study protocol. Resting-state EEG was first acquired, followed by recording during performance of the serial reaction time task (SRTT). Each block of the SRTT included 10 second practice and rest periods across a total of 24 blocks (i.e., 480 seconds of total practice). ***B,*** The mean learning trajectory (average correct keypresses per second for each block) across all participants on the SRTT. Early learning (i.e., where 95% of the learning took place) was determined to occur across blocks 1-10*. **C,*** Online learning refers to learning gains that occur during active practice, i.e., between the beginning and end of a task block. Offline learning refers to learning gains that occur during rest, i.e., between the end of one task block and beginning of the next.

Motor performance was characterised by examining a change in tapping speed (i.e., keypresses made in response to visual cues). Time intervals between correct key presses were calculated in milliseconds (i.e., inter-tap intervals), and then divided by 1000 to derive a measure of key presses made per second. An exponential function was used to model the learning trajectory at the group level (i.e., average increase in correct keypresses per second across blocks) to determine the early learning phase over which period 95% of total learning occurs (as per Bönstrup et al., 2019, 2020; Brooks et al., 2024) (Figure 1b). In this study, the 95% early learning phase occurred across blocks 1-10. As per previous studies investigating micro-consolidation of motor sequences (e.g., Brooks et al., 2024), micro-online learning was defined as the difference in tapping speed between the first and last four correct keypresses of each block. Micro-offline learning was defined as the difference in tapping speed between the last four correct keypresses of a practice block and the first four correct keypresses of the subsequent block (i.e., the difference between the end of block n and the start of block n + 1). Measures of micro-online and offline learning were summed across blocks 1-10 in the early learning phase (Bönstrup et al., 2019).

#### EEG pre-processing

EEG data was pre-processed using the RELAX pipeline (v2.0.0) implemented in the Matlab-based toolbox EEGLAB (v2023.0; Bailey et al., 2023; Delorme & Makeig, 2004). Data were bandpass filtered (0.25 Hz high cutoff, 120 Hz low cutoff) and subsequently downsampled to 500 Hz.The ZaplinePlus function was used to remove 50 Hz line noise and the multi-channel Wiener filter (MWF) was applied to remove muscle and blink artifacts. Independent component analysis (PICARD) followed by ICLabel classification and wavelet-enhanced ICA was used to identify and correct residual artifacts. Rejected channels were interpolated and data was re-referenced to the average. Cleaning quality metrics (signal-to-error ratio, artefact-to-residual ratio, and artifact proportions) were inspected to confirm effective artifact attenuation.

### Statistical analysis

#### Behaviour

To investigate learning outcomes following practice on the SRTT, one-sample *t*-tests were conducted on each outcome variables (micro-online, micro-offline, and total early learning), against a criterion value of zero. Pearson’s correlation examined the between-subjects relationship between micro-online and micro-offline learning.

#### Spectral analysis

For each participant, pre-processed resting-state and SRTT data was imported into Brainstorm (2024). All data was then transformed from the time domain using a series of Morlet wavelets (3 cycle duration), starting at a centre frequency of 1 Hz, increasing to 70 Hz in 1 Hz increments. Power was averaged across specific frequency bands: mu (8–12 Hz), low-beta (13–17 Hz), high-beta (18–29 Hz), low-gamma (30–48 Hz), and high-gamma (52–70 Hz), to generate Spectro-temporal representations across task practice and rest epochs (Figure 2). The 3-minute resting-state data for each participant was trimmed to the middle 2 minutes, excluding the initial and final 30s segments to minimise muscle artifacts and attentional drift.

**Figure 2.**
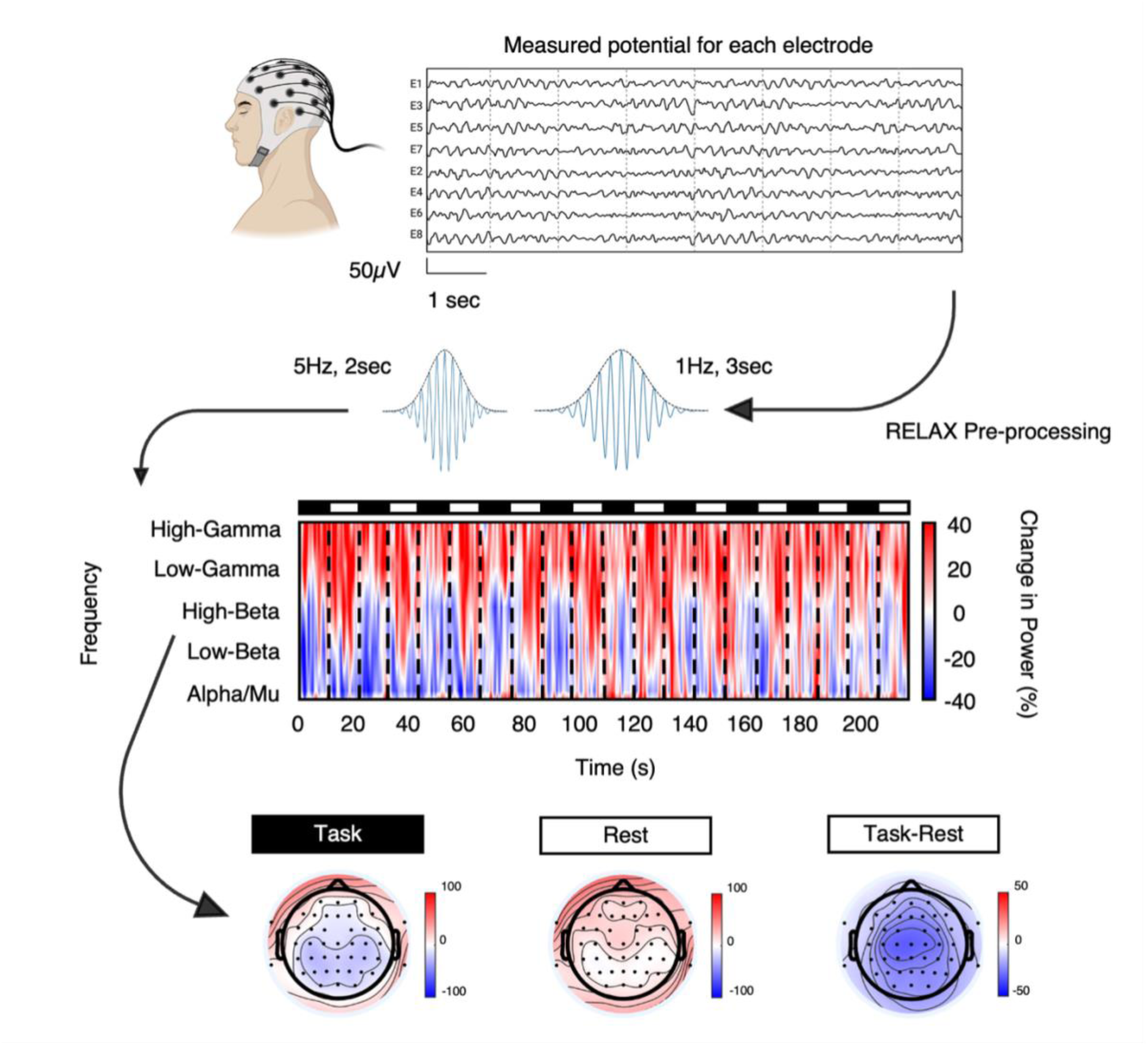
Overview of the analytical approach. EEG was recorded in the time-domain during performance of the SRTT. Following pre-processing, data were transformed by applying a series of Morlet wavelets to generate spectro-temporal representations across specific frequency bands. Percentage change in spectral power was calculated relative to each individual’s pre-task resting-state baseline data, allowing quantification of spectral ERD/ERS.

To ensure comparability with previous research (Bönstrup et al., 2019), the task-related EEG data was normalised to the baseline resting-state data using the following formula:

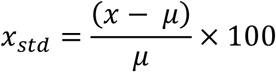

where *x* denotes individual task-related data and *μ* is the individual average (pre-task) resting-state power. The result, *x_std_*, gives a percentage change in spectral power from the pre-task resting-state baseline mean, allowing assessment of synchronisation/ desynchronisation relative to baseline. Percentage change in spectral power was calculated and averaged over each task and rest period per participant for each frequency band of interest (Figure 2).

#### Cluster-based permutation analysis

Cluster-based permutation analyses were conducted using the MATLAB-based toolbox FieldTrip (Oostenveld et al., 2011). Paired-sample *t*-tests examined within-subject differences in EEG spectral power between task and rest conditions for each frequency band (mu, low-beta, high-beta, and the average of low-and high-gamma, hereinafter referred to as gamma). Monte Carlo simulations with 10,000 permutations were conducted (two-tailed), requiring a minimum of two neighbouring electrodes per cluster. The threshold for significance was set by an alpha level of.001, and Bonferroni correction for 63 electrodes and 7 frequency bands (i.e., (0.001/63) / 7 = 2.2676e-06). Statistical results were visualised as topographical maps of *t*-statistics across electrodes.

To assess relationships between EEG and behavioural measures, cluster-based permutation correlation analyses were performed. Two-tailed Spearman’s rank correlations were conducted to assess the relationship between learning measures (micro-online, micro-offline, and total early learning) and oscillatory power at each electrode for each frequency band of interest (mu, low-beta, high-beta, and gamma). Analyses were run separately for task and rest epochs data, and for the difference between task-rest conditions (maxsum cluster statistic, α = 0.05; 10,000 randomisations, requiring a minimum of two neighbouring electrodes per cluster).

## Results

Of the 72 participants that completed the study, 7 participants were excluded from analyses due to potentially having explicit awareness of the repeating motor sequence (i.e., accurately reproducing at least the first 5 items of the 12-item sequence from memory after completing the SRTT). Additionally, 1 participant was excluded from the behavioural analyses, and 7 participants were excluded from EEG analyses due to noisy (e.g., excessive muscle contribution in EEG) and/or missing data. As a result, a total of 64 participants were included in the behavioural data analyses, and 57 participants were included in the EEG analyses. For correlation analyses, only participants with both EEG and behavioural data were included (*n* = 57).

### Micro-consolidation occurs during early implicit sequence learning

Significant learning was observed across the early learning phase of the SRTT (*M* = 0.62, *SD* = 0.26) [*t*(63) = 19.24, *p* <.001, Cohen’s *d* = 2.35, 95% CI 1.92, 2.89] (Figure 3a). This learning was primarily attributable to performance improvements during micro-offline periods, with a significant micro-offline effect (*M* = 0.59, *SD* = 2.12) [*t*(63) = 2.24, *p* =.029, Cohen’s *d* = 0.64, 95% CI 0.03, 0.53]. In contrast, there was no significant micro-online effect (*M* = 0.28, *SD* = 2.16) [*t*(63) = 1.02, *p* =.312, 95% CI-0.12, 0.37] (Figure 3b).

**Figure 3.**
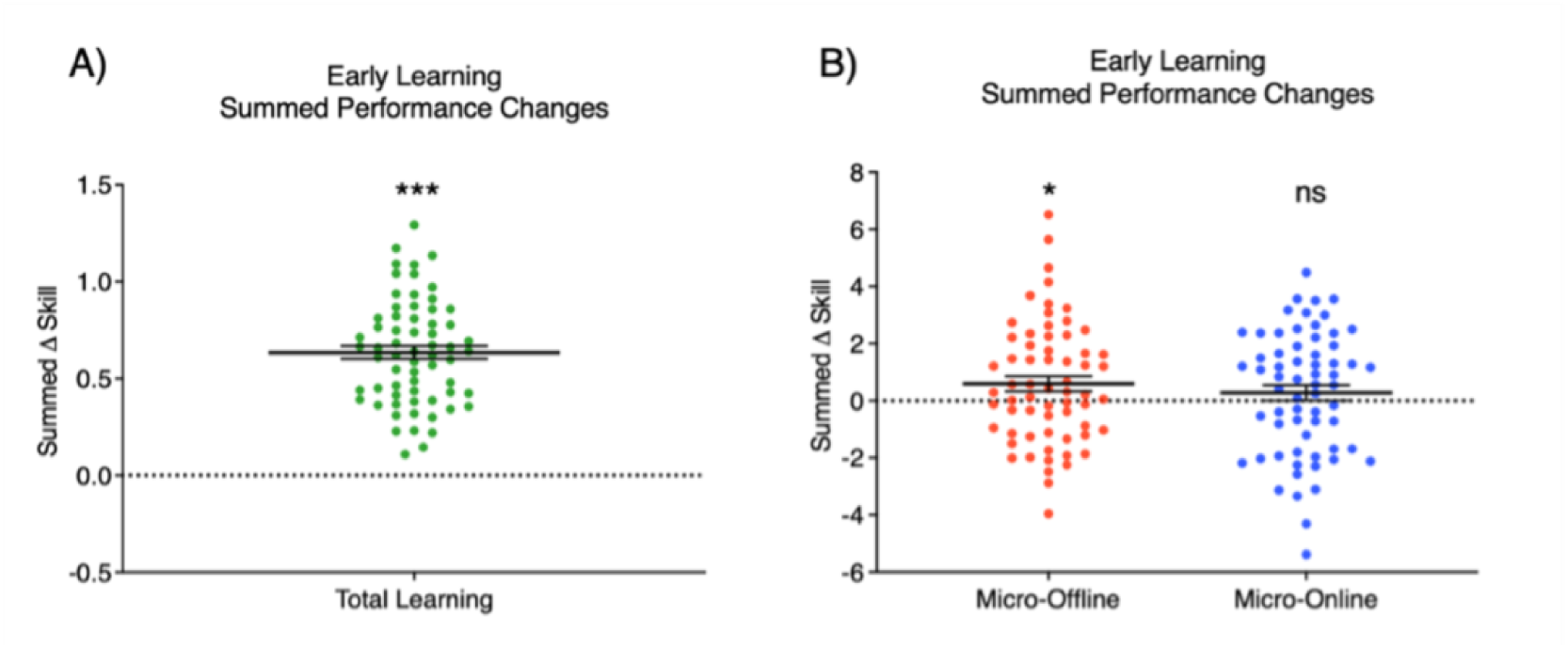
Total early learning (***A,*** green colouration) on the SRTT is primarily attributable to micro-offline gains (***B,*** red colouration). No significant changes were observed during practice (i.e., micro-online periods) (***B,*** blue colouration). Ns nonsignificant, * p <.05, ** p <.01, *** p <.001.

### Modulation of mu and beta power during learning

To examine modulation of neural oscillations during learning, changes in event related spectral perturbations (ERSPs) were assessed between task and rest epochs of the SRTT. Significant ERSPs were observed between task and rest periods for the low-and high-beta bands, as well as for alpha at posterior electrodes (Figure 4d). Specifically, there was a widespread significant reduction in task epochs relative to rest for both low-beta (*M_Task_* = - 10.30, *SD* = 48.13; *M_Rest_* = 11.15, *SD* = 51.11; *t*(56) =-8.45, *p* <.001, Cohen’s *d* = 1.12, 95% CI-1.45,-0.78), and high beta (*M_Task_* =-9.47, *SD* = 47.09; *M_Rest_* = 22.46, *SD* = 50.42; *t*(56) =-13.91, *p* <.001, Cohen’s *d* = 1.84, 95% CI-2.27,-1.41) (Figure 4a-d). No significant differences were observed in the sensorimotor mu and gamma bands (all p >.39) (Figure 4d). To further visualise this effect, a time-frequency plot of the group mean ERSP values in each frequency band of the SRTT is shown for electrode Cz in Figure 2, and per block averaged across all significant electrodes in the cluster in Figure 4e. On average, a suppression in beta band power is evident following the onset of the 10s task practice periods (i.e., ‘online’ learning), followed by comparatively higher beta band power during the offline 10s rest periods, during which time micro-offline learning occurs.

**Figure 4.**
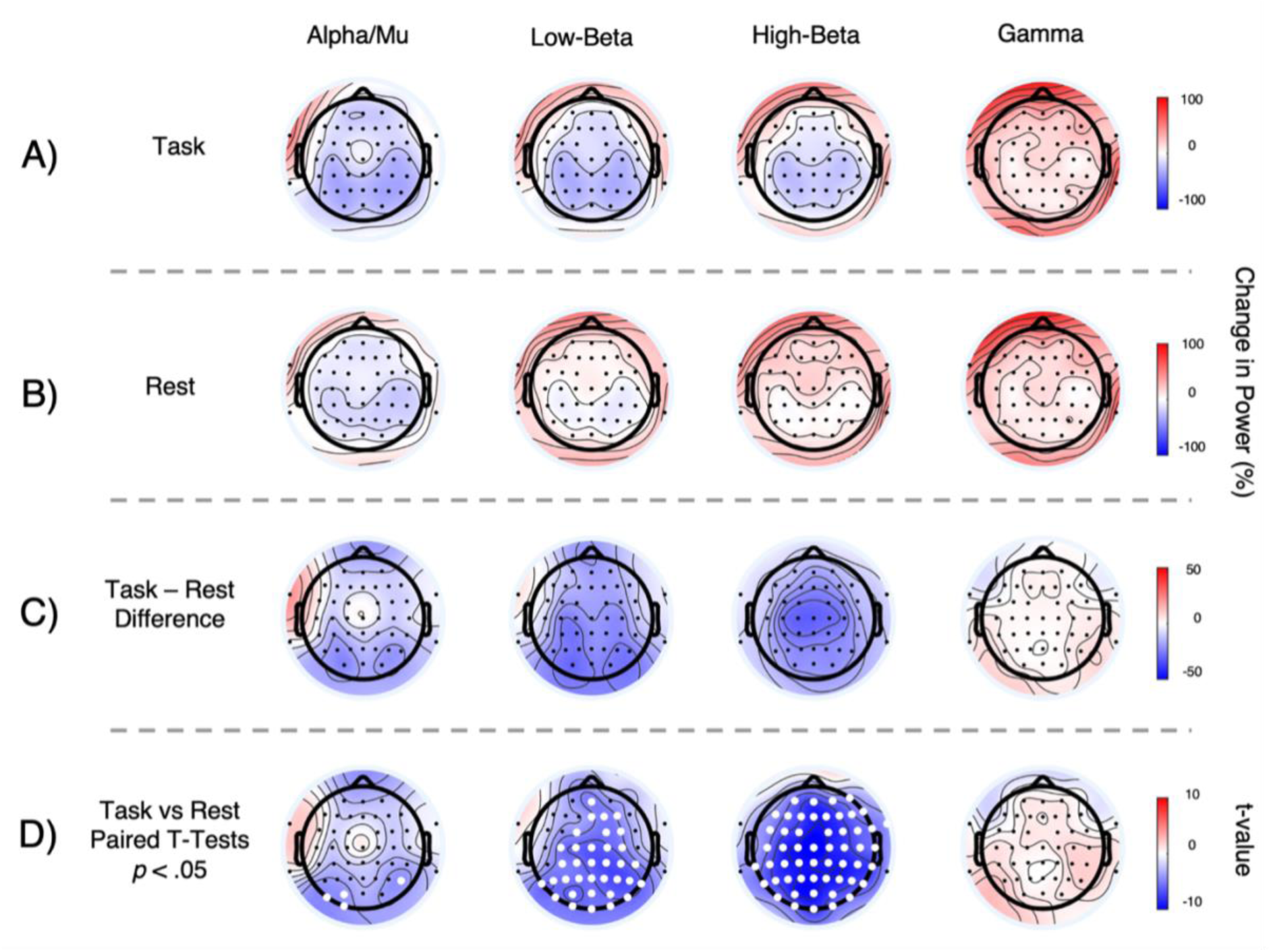
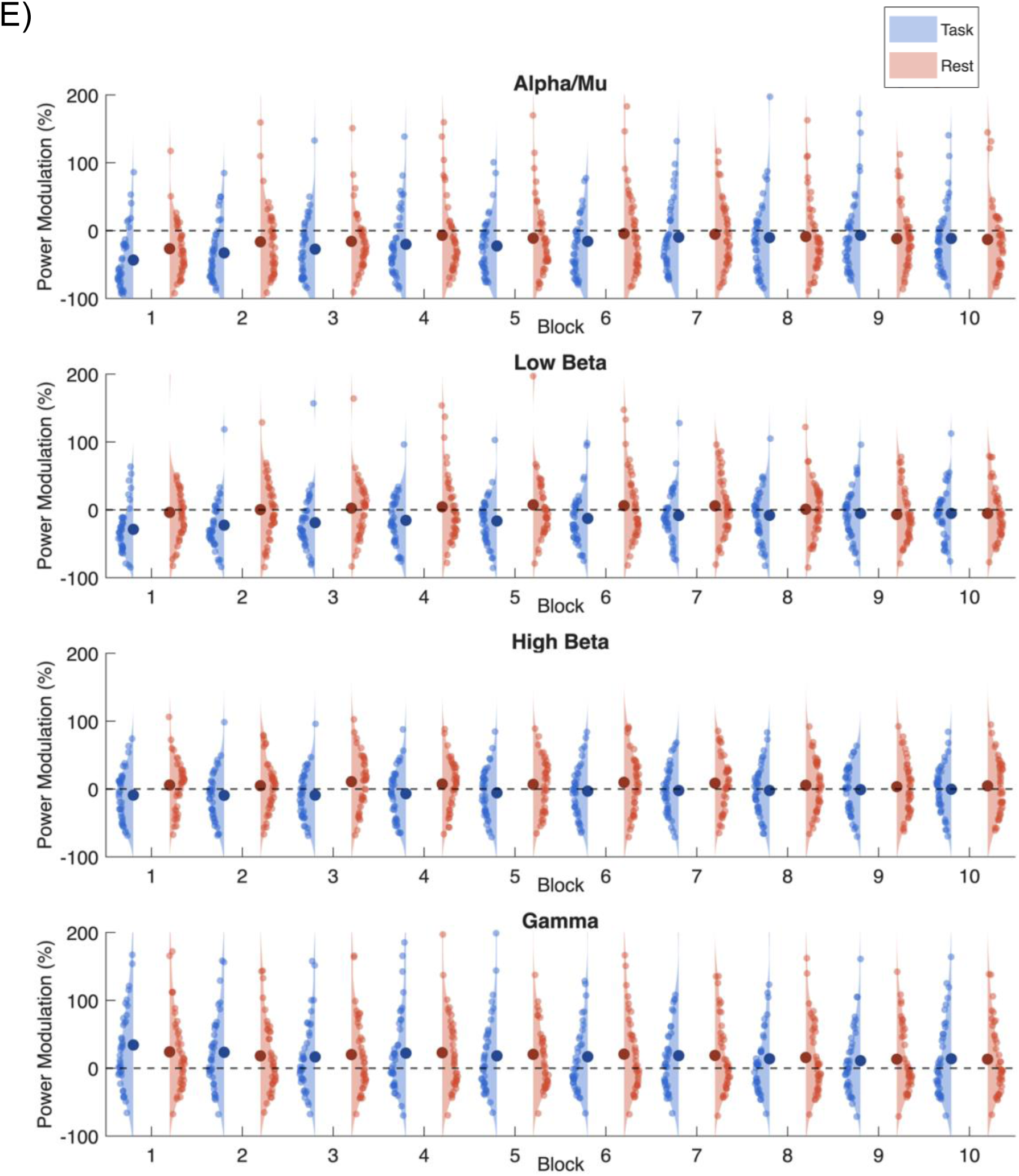
***A-C,*** Spectral power change across task **(A)** and rest **(B)** blocks of the SRTT. A significant reduction (depicted in blue colouration) in beta power was observed during practice relative to rest **(C)**, across a distributed topography **(D)** with significant electrodes depicted in white. This effect was specific to beta and was not observed across mu or gamma rhythms. **(E)** Plots depicting average change in oscillatory power across task (blue) and rest blocks (red) across the early learning period. Violin plots with individual subject values depict the distribution. Larger bolded circles represent the mean for mu, low-beta, high-beta, and gamma bands.

### Micro-consolidation of implicit motor sequences is associated with beta power modulation

Cluster-based permutation correlation analyses evaluated relationships between learning and oscillatory power. Greater micro-offline gains were associated with reduced high-beta power during the rest epochs of the task [*r*_s_(56) =-.33, *p* =.011, 95% CI-0.55,-0.07] across a frontal-central topography (Figure 5). This relationship was specific to high-beta (p >.05 for all other frequency bands).

**Figure 5.**
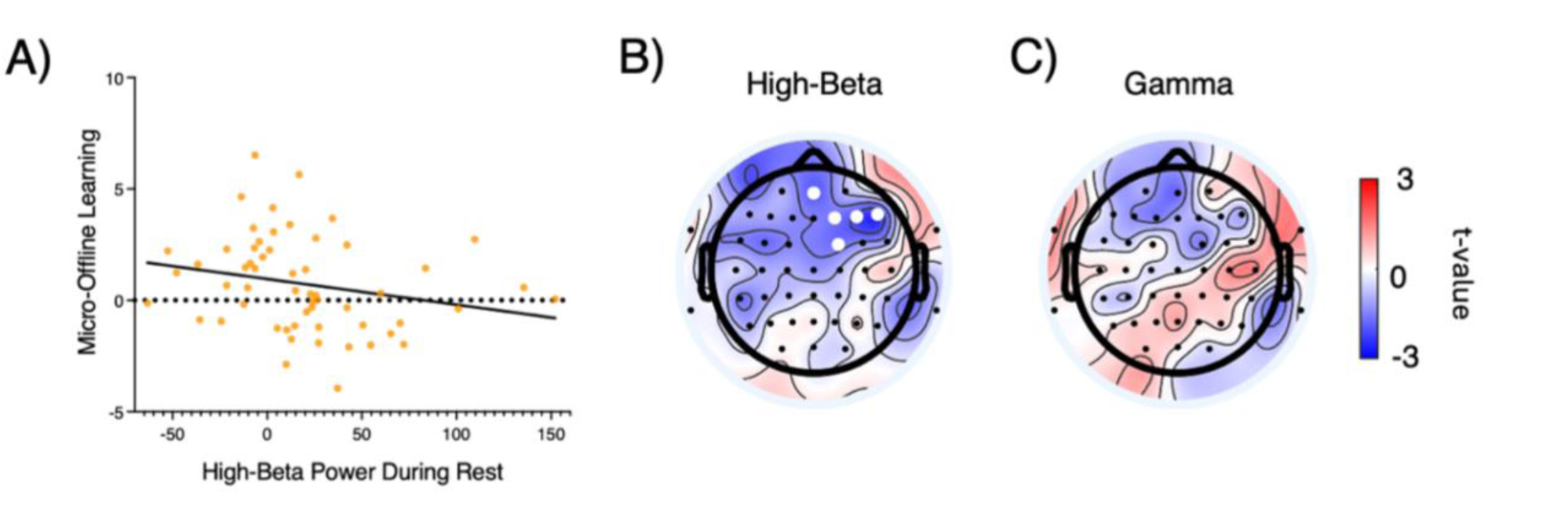
**(*A)*** Micro-consolidation is associated with modulation of beta power across a frontocentral topography. The x-axis shows high-beta power from the resting epochs of the micro-consolidation phase of the task, expressed as a percentage of a pre-task resting state EEG, see methods for further detail. Across participants, a reduction in high-beta power was associated with a greater micro-offline learning effect (B). This relationship was specific to high-beta and was not observed for the gamma frequency band (C).

## Discussion

In this study, we investigated the neural oscillations supporting micro-consolidation of an implicit motor sequence. Here, we reproduce our previous finding of micro-consolidation during the early stages of implicit sequence learning (Brooks et al., 2024) in an independent sample twice the size. We show that the early ‘fast’ learning on an implicit sequence task is primarily expressed as micro-offline gains during brief rest periods between periods of practice. We then show consistent modulation of beta power as a function of active practice vs rest during implicit learning, and demonstrate that micro-offline gains are associated with high-beta power during the rest epochs. Specifically, weaker high-beta power during the rest epochs was associated with greater micro-offline gains. This relationship was observed at frontocentral electrodes, and was specific to the beta frequency, as relationships were not observed for either the mu or gamma bands. Overall, in line with seminal work implicating beta in early explicit motor learning (Bönstrup et al., 2019; Buch et al., 2021), our results indicate that beta is a shared neurophysiological signature of micro-consolidation of implicit sequences.

In line with our expectations, we observed significant micro-offline improvement during the early learning of an implicit motor sequence. This finding is consistent with both our recent work (Brooks et al., 2024) and a growing body of research across explicit and probabilistic learning paradigms reporting rapid offline consolidation during early ‘fast’ phase of skill learning (see Brooks et al. (2026) for review). Collectively, our results, in conjunction with existing evidence across distinct motor learning paradigms (e.g., explicit and higher-order probabilistic learning tasks) (Brooks et al., 2026), demonstrate that early skill learning benefits from brief rest periods which facilitate rapid offline consolidation of skill.

Previous work has linked micro-consolidation of explicit sequence learning to frontoparietal beta oscillations assessed with magnetoencephalography (Bönstrup et al., 2019). Here we show, using EEG, that desynchronisation of frontocentral beta rhythms delineate periods of practice and rest during the early stages of implicit skill learning. Our findings extend upon previous work implicating transient beta oscillations in motor skill error processing and adaptation (Koelewijn et al., 2008; Tan et al., 2014; Torrecillos et al., 2018) and in stabilisation and consolidation of both uni-and bimanual skills (Pfurtscheller & Lopes da Silva, 1999; Pollok et al., 2014; Wu et al., 2025). While some studies report transient (e.g., 1s) event-related synchronisation of beta power following movement cessation that may signal error-based learning (e.g., Koelewijn et al., 2008; Wu et al., 2025), here we demonstrate modulation of beta power across comparatively extended time windows, with an average reduction in beta over 10s practice periods, followed by a relative increase over successive 10s periods of rest (albeit still reduced compared to the pre-task resting state in many participants, see Figure 5). Collectively, these findings add to a growing literature implicating beta in distinct features of skill acquisition, from error monitoring and adaptation to the rapid consolidation processes supporting early learning.

We also demonstrate that the degree of beta power during the 10 s rest epochs is associated with the magnitude of micro-offline gains. While the functional role of beta rhythms in micro-consolidation is yet to be clearly established, we reason that the relationship between beta and micro-offline improvement may reflect the reduction of inhibition which occurs during early learning (Coxon et al., 2014). Several studies have associated beta ERS with GABA-mediated disinhibition across task-relevant brain networks (Gaetz et al., 2011; Hall et al., 2011; Muthukumaraswamy et al., 2013), and disinhibition is a prerequisite for the plasticity supporting memory consolidation (Chen et al., 2015; Stagg et al., 2011). This framework aligns with our observation of greater ERD during practice and weaker beta ERS over the 10 s rest epochs of the early ‘fast’ phase of learning, which then recovers with time and practice (see also Wu et al., 2025). Indeed, recent work including our own has implicated sensorimotor GABA concentration across the early stages of skill learning when micro-consolidation processes are orchestrated (Hendrikse et al., 2025; Kolasinski et al., 2019). Our finding that weaker beta is associated with higher micro-offline learning establishes a role of beta oscillations that generalises across rapid consolidation of both explicit and implicit forms learning (Bönstrup et al., 2019; Griffin et al., 2025). Notably, we show that relationship between micro-consolidation of implicit motor learning is specific to high beta (18-29 Hz). Broadly, this aligns with previous work linking reduced frontoparietal beta (16-22 Hz) to micro-consolidation of explicit learning (Bönstrup et al., 2019), and showing a reduction of micro-consolidation following 20 Hz beta alternating current stimulation (Griffin et al., 2025). Future work is required to delineate the functional contribution of low vs high beta and establish the interplay between beta, GABA-mediated inhibition, and rapid skill consolidation.

Reduced beta power may also facilitate aspects of systems-level replay. Across the literature to date, there is evidence linking micro-consolidation of explicit motor sequences to sharp wave ripples and neural replay processes (Bönstrup et al., 2019; Griffin et al., 2025; Jacobacci et al., 2020). For instance, intracranial electrode recordings from macaque monkeys have linked an increase in cortical ripples to micro-offline gains in performance of a visuomotor sequence (Griffin et al., 2025). Intriguingly, this micro-offline improvement was also associated with a reduction in beta bursts (Griffin et al., 2025), plausibly reflecting a reduction of GABA-mediated inhibition (Coxon et al., 2014; Hendrikse et al., 2025). Magnetoencephalography work in humans has also associated micro-consolidation of explicit sequences with neural replay across cortico-hippocampal networks, which involves a temporal compression of the learnt behaviour within the beta frequency (Buch et al., 2021). In this context, involvement of beta may indicate a removal of inhibition, providing a window for the encoding, reactivation, and maintenance of the learnt sequence and task-relevant information across cortical-hippocampal networks (Brooks et al., 2026).

### Future Directions and conclusion

Our findings should be considered in the context of certain limitations. For instance, our findings establishing beta modulation as a signature of implicit micro-consolidation are inherently correlational. Future work aimed at establishing causal relationships via application of perturbation techniques (e.g., non-invasive brain stimulation) will be informative in establishing the functional role of beta modulation in early skill learning in humans. One approach may be to apply transcranial alternating current stimulation at the beta frequency across task-relevant networks (e.g., frontocentral cortical regions implicated in sensorimotor control) during early stages of skill learning (Joundi et al., 2012; Leunissen et al., 2022), or conversely apply forms of stimulation capable of directly targeting deep brain structures implicated in micro-consolidation such as the hippocampus (Brooks et al., 2026). By extension, establishing how modulation of early learning processes (e.g., via causally perturbing task-relevant structures) influences beta power and motor skill across longer timescales (e.g., days-weeks) is also an outstanding question which warrants further investigation.

In conclusion, we provide evidence implicating a state of reduced beta power during rest epochs in the micro-consolidation of an implicit motor skill. Our findings add to a growing body of work highlighting the significance of rapid consolidation processes during the early ‘fast’ stages of skill learning across distinct domains of motor performance. Establishing causal links between beta and micro-consolidation may inform novel approaches to optimise motor learning and performance across health and disease.

## Additional information

### Author contributions

J.H., N.A., and J.C. were involved in the study design. J.H., N.A., J.H., E.S., E.M., S.T., and J.C. collected the data. J.H., N.A., J.H., E.S., E.M., S.T., and J.C. were involved in the analysis and interpretation of the data. J.H., N.A., J.H., E.S., E.M., S.T., and J.C. contributed to writing and/or revising the manuscript. All authors approved the submitted manuscript, and agree to be accountable for all aspects of the work in ensuring that questions related to the accuracy or integrity of any part of the work are appropriately investigated and resolved. All persons designated as authors qualify for authorship, and all those who qualify for authorship are listed.

## Acknowledgements

The authors would like to thank the participants who volunteered their time for this research. Open access publishing facilitated by Monash University, as part of the Wiley - Monash University agreement via the Council of Australian University Librarians.

## Data availability

De-identified behavioural and EEG data are available at https://osf.io/guzs8/. Readers seeking access to the data and/or task materials should contact the lead author. All code used for cognitive and MRI analysis is available at https://github.com/jhendrikse/.

